# Neuroinflammation induces nerve growth factor dependent nociceptor sensitisation in a neonatal rodent model of platinum-based chemotherapy induced neuropathic pain

**DOI:** 10.1101/2023.09.21.558866

**Authors:** Marlene Da Vitoria Lobo, Lydia Hardowar, Tameille Valentine, Lucy Tomblin, Charlotte Guest, Dhyana Sharma, Mark Paul-Clark, Richard Philip Hulse

**Affiliations:** Department of Biosciences, School of Science and Technology, Nottingham Trent University, Nottingham, NG11 8NS; Division of Cancer and Stem Cells, School of Medicine, University of Nottingham, Nottingham NG7 2UH

**Keywords:** nociceptor, dorsal root ganglia, NGF, chemotherapy, cisplatin, neurodegeneration, TRPV1, inflammation, macrophage

## Abstract

Chemotherapy-induced neuropathic pain (CINP) is a common adverse health related comorbidity that manifests later in life in paediatric patients treated for cancer. CIPN pathology progressively develops over time resulting in a delayed but long-lasting neuropathic pain. Current analgesic strategies are ineffective, aligning closely with our lack of understanding of CINP. Recent studies have indicated alterations in sensory neuronal maturation as component of CINP. The aim of this study was to investigate how cisplatin induces nerve growth factor mediated neuroinflammation and nociceptor sensitisation. In a rodent model of cisplatin induced survivorship pain, there was a significant infiltration of nerve growth factor positive macrophages into the dorsal root ganglia (DRG), demonstrating a robust neuroinflammatory response. Additionally, it was observed that CD11b/F480 positive monocyte/macrophages challenged with cisplatin expressed more NGF. Additionally, DRG derived primary sensory neuron cultures from neonatal mice demonstrated enhanced NGF-dependent TRPV1 mediated nociceptor activity after cisplatin treatment. Increased nociceptor activity was also observed when cultured neurons were treated with conditioned media from cisplatin activated monocyte/macrophages. This elevated nociceptor activity was dose-dependently inhibited by a neutralising monoclonal antibody to NGF. Intraperitoneal administration of NGF neutralising antibody significant reduction in mechanical hypersensitivity was given to mice with cisplatin-induced juvenile survivorship pain there was a as well as suppression of cisplatin induced aberrant nociceptor intraepidermal nerve fibre density. These findings identify the NGF/TrkA signalling pathway as a potential novel therapeutic target for analgesia in adult survivors of childhood cancer.

## Introduction

Current advances in medical advances in diagnosis and treatment for paediatric cancer, has led to significant increases in survival rates [16]. Unfortunately, due to the intensive and invasive nature of the adopted chemotherapeutic approaches, patients suffer from extensive impact upon their quality of life [16]. This is highlighted by adverse health related implications such as chronic pain, nephrotoxicity, optotoxicity, fatigue and negative emotive cognitive disorders as well as associated societal impact. Persistent long-lasting pain in childhood cancer survivors is prevalent in upto 59% of individuals and is attributed to the intensive nature of treatment [1,12]. Pain frequently develops in adolescence and persists into adulthood well beyond diagnosis and cessation of treatment [15]. Unfortunately, there are no condition-tailored analgesics available for child cancer survivorship pain as analgesia is predominantly ineffective and/or causes adverse health related side effects in the long-term [2]. These difficulties arise due to analgesic management being based upon other neuropathic pain conditions and a complete lack of understanding to what causes chronic pain in ASCC.

A front-line cancer treatment is platinum-based chemotherapy, which is effective against a range of solid tumours, including lung, bladder, testicular, ovarian cancers and metastatic testicular germ cell cancer [7]. However, there are significant limitations to the use of these chemotherapy agents due to the severe, dose-limiting side effects of platinum-based chemotherapy treatments. A major dose-limiting, and potential cause for premature termination of treatment, is the neurotoxic effect of platinum-based chemotherapeutics on the peripheral somatosensory nervous system [14]. This cumulative sensory neuronal damage caused by chemotherapy treatment has been extensively reported to induce both immediate and delayed-onset pain sensations, that can manifest years after the cessation of treatment, well into adulthood [11,14,15]. It is important to note that the somatosensory nervous system is vulnerable at an early age to exposure to cellular stress/damage[6,19]. When the somatosensory nervous system is exposed to a traumatic injury or inflammatory insult during neonatal development, pain hypersensitivity develops but is delayed with onset occurring during adolescence and persisting throughout adulthood [5]. The development of this delayed nociceptive behavioural phenotype is thought to arise in part from adaptations in the inflammatory system. Induced proinflammatory events are typically impaired until later in life, but are unmasked during an insult early in life [13]. We have developed a rodent model to allow investigation into platinum-based chemotherapy induced childhood cancer survivorship pain [9]. This enables exploration into the causative mechanisms that drives the onset of this chronic pain state and determine efficacy of potential analgesics. In these studies we demonstrate that cisplatin induced pain that occurs later life is NGF dependent, mediated via a pro-neuroinflammatory state within the dorsal root ganglia.

## Methods

### Ethical Approval and Animals used

All experimental procedures involving animals were carried out in consultation with local Animal Welfare and Ethics Review Board (Nottingham Trent University) and in accordance with UK Home office animals (Scientific procedures) Act 1986 and ARRIVE guidelines. Animals had ad libitum access to standard chow and were housed under 12:12h light:dark conditions.

### Induction of Cisplatin induced neuropathic pain and administration of pharmacological agents

Neonatal Wistar rats were administered either vehicle (phosphate buffered saline; PBS) or cisplatin (Sigma-Aldrich; 0.3mg/kg) via intraperitoneal injection delivered on 2 occasions; postnatal (p) day 14 and 16. All rodents were weaned no later than 22 days and group housed according to gender. Males and female Wistar rats were utilised in all outlined studies, with animal number outlined in figures. Body weight was monitored regularly throughout the study, with no observed reductions in body weight identified.

### Nociceptive Behaviour

All rodents were habituated to the testing environment prior to nociceptive behavioural experimentation [3,22]. Mechanical withdrawal thresholds were acquired following application of Von Frey (vF) hairs to the plantar surface of the hindpaw. Withdrawal thresholds were calculated following application of differing vF filaments of increasing force, with each vF applied a total of five times to the plantar surface of the hindpaw. Force response withdrawal curves were generated and mechanical withdrawal thresholds were determined as the mechanically applied force to elicit 50% of nociceptive withdrawals. The Hargreaves test was performed to determine heat nociceptive withdrawal latency [8]. A radiant heat source was applied to the plantar surface of the hindpaw. The duration (latency) between onset of stimulus to the mouse withdrawing their paws was recorded as the withdrawal latency. This was measured 3 times and a mean latency was calculated for both hindpaws.

### Drug Delivery

1μM Nerve Growth Factor 2.5s (NGF, Alomone) was administered via a 20μl subcutaneous injection under recovery anaesthesia (∼2% isoflurane in O_2_). In some instances, NGF neutralising antibody was administered via intraperitoneal injection (0.1mg/kg) or 2μg/ml via subcutaneous injection in phosphate buffered saline versus IgG control.

### Immunofluorescence

Animals from each experimental group were terminally anaesthetised (Dolethal; intraperitoneal sodium pentobarbital 200mg/ml) and were transcardially perfused with PBS and subsequently by 4% paraformaldehyde (PFA; pH7.4). Lumbar dorsal root ganglia (DRG) and hindpaw plantar skin were extracted, submerged in PFA overnight and cryopreserved in 30% sucrose. Tissues were stored at -80oC until processing [20]. Cryosections for DRG (8µm thickness) and plantar skin (20 µm thickness) sections were collected onto superfrost plus slides (VWR International). Slides were washed 3 times with PBS 0.2% Triton and then blocked with PBS, 0.2% Triton, 5% BSA (bovine serum albumin), and 10% FBS (foetal bovine serum) for 1 hour at room temperature. Sections were incubated at 4°C for 72 hours in blocking solution (5% bovine serum albumin, 0.2% Triton-X) containing primary antibodies CD45 antibody (Abcam, ab10558, 1:400 dilution), CD11b (Abcam, 1:200), anti-NGF (Sigma Aldrich, N8773 1:100), IB_4_ biotinylated (Sigma Aldrich, L2140, 1:200). Sections were washed in PBS for 5 minutes three times. Secondary antibodies were subsequently applied to the tissue sections in PBS + 0.2% Triton X-100 at room temperature for 2 hours. CGRP immunoreactivity required an additional step to incorporate anti-rabbit biotinylated IgG (1 in 500, Jackson Laboratories 2hrs at room temperature) following by streptavidin alexafluor. Secondary antibodies (1:1000, all Invitrogen, UK) used were Alexa Fluor 555-conjugated donkey anti-mouse (ab150114), Alexa Fluor 555-conjugated donkey anti-rabbit (ab15158), Alexa Fluor 488-conjugated donkey anti-rabbit (ab150073), and streptavidin-conjugated Alexa Fluor-555 (S32355). Coverslips were mounted using vector laboratories H1000 medium set mounting medium. Sections were imaged via confocal microscopy using a Leica confocal microscope.

### Endothelial Cell Culture

Human umbilical vein endothelial cells (PromoCell C-12203) (HUVEC) were cultured and plated in either 6, 24 or 96 well plates. For Western blot analysis 50,000 cells per well were seeded on two 6 well plates, 2,000 cells per well were seeded in a 96 well plate for viability assays and 5,000 cells per well were seeded in a 24 well plate with ethanol sterilized coverslips for immunofluorescence assays. Cells were left to grow until they reach 90% confluency and then treated with cisplatin (0, 1, 3 or 5 μg/ml) for 24 hours.

### Immunocytochemistry

Coverslips from the 24 well plate were fixed with 1% PFA for 10 minutes and subsequently washed 3 times with PBS 0.1% BSA, followed by one wash of PBS 0.1% Tween, 3 more PBS/BSA washes and then blocked using the blocking solution. Primary antibodies for VE-cadherin (Abcam, ab33168, 1:100) or ICAM-1 (Santa Cruz, 1:200) were added and left overnight at 4 °C. Three washes with PBS/BSA were performed after the primary antibodies incubation. Secondary antibodies (anti rabbit Alexafluor488 or 555) were added in a 1:500 dilution for 1 hour at room temperature. Three more washes were performed and the coverslips were carefully removed from the 24 well plate, placed onto Super Frost slides (Sigma Aldrich) and mounted with a medium set VectaShield (H1000). Coverslips were sealed with nail polish. All images were taken using the LAS X software from Leica Microsystems for confocal microscopy (TCS SPE confocal microscope).

### Splenocyte adherence assay

Spleens were removed from adult C57bl6 mice and were passed through a 40μm strainer using RPMI 1640 medium (Invitrogen), penicillin-streptomycin (Sigma), 10% foetal bovine serum (Invitrogen), l-glutamine (if not already in RPMI) (Invitrogen), sodium pyruvate solution (Sigma) amd monothioglycerol (Sigma). Cells were incubated in Red Blood Cell Lysing Buffer (R7757 – Sigma) prior to experimentation. Splenocytes from were labelled with Vybrant Dil before incubation with cisplatin-treated HUVEC. Splenocytes in suspension were incubated at 37 °C, 5% CO_2_ with 5 μl of Vybrant Dil solution per ml of suspension for 20 minutes. Media was removed by centrifugation and the washing procedure was repeated twice. Splenocytes were incubated with cisplatin treated HUVECs for 24 h in a 24 well plate. The cells were then fixed with 1% PFA for 10 minutes and subsequently washed 5 times with PBS. Coverslips were removed from the plate, placed onto Super Frost slides using medium set Vectashield mounting medium and finally sealed with nail polish for confocal microscopy imaging.

### Western Blot

Endothelial Cells; Protein from treated HUVEC was extracted using RIPA buffer (ThermoFisher) with 1X protease inhibitor cocktail (ThermoFisher, 78440). Protein lysate was equally loaded onto 10% Precast gels (Bio Rad), with 50 μg of protein per well. Membranes were blocked with TBS 1% BSA, 0.1% Tween for 1 hour at 4 °C. Primary antibodies were used in a 1:200 dilution for ICAM-1 and β-actin, with 1 μg/ml for VE-cadherin and occluding utilised. Primary antibodies were left overnight at 4 °C. Three washes with TBS, 0.1% Tween were done after the incubation. Secondary Licor antibodies were used in a 1:5000 dilution and were incubated for 1 h at room temperature. Five final washes with TBS, 0.1% Tween were performed and membranes were then analysed with the Licor Odissey.

### Splenocytes

Protein lysate samples were extracted from mouse splenocytes. Splenocytes were plated into 6 well plates for 24h prior to cisplatin (5ug/ml) treatment. Cells were lysed using RIPA buffer (Sigma-Aldrich) containing protease inhibitor cocktail (Protease/Phosphastase inhibitor cocktail, Cell Signalling). Equal protein lysate concentrations (40μg per well) were loaded on a 4%-20% precast Mini-Protean gradient TGX gel (Biorad). Proteins were separated by SDS-PAGE electrophoresis and transferred to PVDF membranes using Trans-blot turbo transfer system (Bio-Rad). Membranes were incubated in 5% BSA in tris-buffered saline (TBS)-Tween 0.1% (TBST) for 1 hour at room temperature. Primary antibodies were incubated overnight at 4 °C (NGF, CD45, B actin). The membrane was then incubated in secondary antibodies (Licor donkey anti-rabbit, anti goat and anti-mouse antibodies 1:10000) in TBST-0.1%-1% BSA. Membranes were washes three times with TBST and visualised on the Licor Odyssey Fc.

### Primary Dorsal Root Ganglia Sensory Neuronal Cell Culture

Prior to dissection, 75μL per well of poly-L-lysine was added to a 96-well plate and incubated overnight. The wells were washed with PBS, left to dry, then 100μL of 5μg/mL laminin in PBS was added to each well. Dorsal root ganglia were extracted from C57BL/6 mouse pups aged between postnatal days 0-7 and collected in Hams/F-12 media, 3% BSA, penicillin/streptomycin and N-2 supplement. Enzymatic digestion was performed using collagenase type IX, with DRGs incubated for 2 hours. This cell suspension was mechanically triturated using a 1mL pipette tip to ensure the complete dissociation of the neurons, with 1mL of the cell suspension added to each of two 15% BSA cushions (1mL Ham’s F12 media + 1mL 30% BSA) in 15mL falcon tubes and centrifuged at 1200rpm for 8 minutes. The supernatants containing the cell debris, BSA and media was aspirated, and the cell pellets containing the neurons were resuspended in 1mL of media. DRG sensory neurons were seeded at a concentration of 2000 cells per well.

### DRG Sensory Neuronal Calcium Assay

Final concentrations of NGF and capsaicin added were 1nM and 1μM respectively. These treatments were added 24 hours prior to imaging. For the cisplatin treatments with NGF and antibody, all treatment groups were treated with 5μg/mL cisplatin made up in supplemented F12 media one week prior to imaging, with media replaced with freshed media 24 hrs later (6 days prior to calcium assay). DRG sensory neurons were loaded with 100μL per well of Fluo-4 cell permeant in media containing 5% Pluronic acid, and incubated for 1 hour. Capsaicin evoked activity was measured by the Infinite M Plex plate reader. The fluorescence response per well was measured at 10 second intervals over a 200 second time period. Prior to imaging, the plate reader was set to measure at 37°C to reflect physiological temperatures. The wavelength settings were selected based on the manufacturer specifications (488nm excitation and 516nm emission), with the gain set to 185. Baseline fluorescence readings were taken prior to each capsaicin treatment as t=0, with the subsequent timepoint readings expressed as a fold change over said baseline.

### Statistical Analysis

All data are represented as mean±SEM unless stated. Data were acquired and quantified using Microsoft Excel 2010, Image J (https://imagej.nih.gov/ij/) [17,18]and Graphpad Prism 8. Raw Ca2+ response values obtained were collated and background fluorescence removed by subtracting the smallest measured fluorescence value from each value in the dataset. These new values were normalised by dividing by the fluorescence value obtained prior to capsaicin addition as a control, in order to determine the fold response over basal levels. For the capsaicin dose-response, a two-way ANOVA was employed to determine the treatment effect over time at different concentrations. fdFor all other data sets, the effects of different treatments on the Ca2+ neuronal response were compared by area under the curve using T-student and one way ANOVA tests were used depending on the data retrieved from each experiment. Nociceptive behavioural assays were determined using one way ANOVA with post Bonferroni test. Immunohistolgical analysis utlised Unpaired T Tests or one way ANOVA with post Bonferroni test.

## Results

Early life administration of cisplatin induced a delayed but pronounced mechanical allodynia (Fig.1A), with no observed alterations in body weight (Fig.1B). In addition, neonatal rodents administered cisplatin during the second week of life demonstrated a significant proinflammatory profile in the peripheral somatosensory nervous system; DRG and plantar skin at a timepoint accompanying presentation of pain (Fig.2). In the plantar skin (age matched vehicle Fig.2 A&B), cisplatin induces an increased infiltration and accumulation of CD45 positive monocyte/macrophage (Fig.2 C&D, G) versus age matched vehicle controls. Similarly, in comparison from the plantar skin of age matched vehicle controls there was an increase in CD11b positive cell number in plantar skin from cisplatin treated rodents (Fig.2E&F, H; No primary control representative image provided Fig. 2I). Additionally, cisplatin also induced infiltration of CD45 positive monocyte/macrophages (Fig. 2J-L) in the lumbar dorsal root ganglia when compared to age matched vehicle controls, as well as activation of resident satellite glia (GFAP positive, Fig.2 M&O). In the plantar surface of the hindpaw of cisplatin treated rodents, there was an increase in CD45 inflammatory cell accumulation in association with microvessel endothelium (IB4 labelled) compared to age matched vehicle controls (Fig. 2P&Q) indicative of enhanced monocyte/macrophage adherence and infiltration. This infiltration and accumulation of CD45 and CD11b cell types in the peripheral somatosensory nervous system, was associated with alterations in endothelial cell function in relation to permeability and cell adhesion (Fig. 3). Increased inflammatory cell-endothelial cell adherence was associated with elevations in ICAM1 expression in cisplatin treated endothelial cells (HUVECs and spinal endothelial cells; Fig. 3A-F). Additionally, cisplatin induced loss of tight junctional proteins (VE-Cadherin Fig. 2G-J and Occludin Fig. 3I&K) promotes increased capillary leakiness and cell trafficking. This was demonstrated in cisplatin treated HUVECs having increased adherence of fluorescently (Dil) labelled splenocytes compared to vehicle treated cells (Fig. 3l&M).

**Figure 1.**
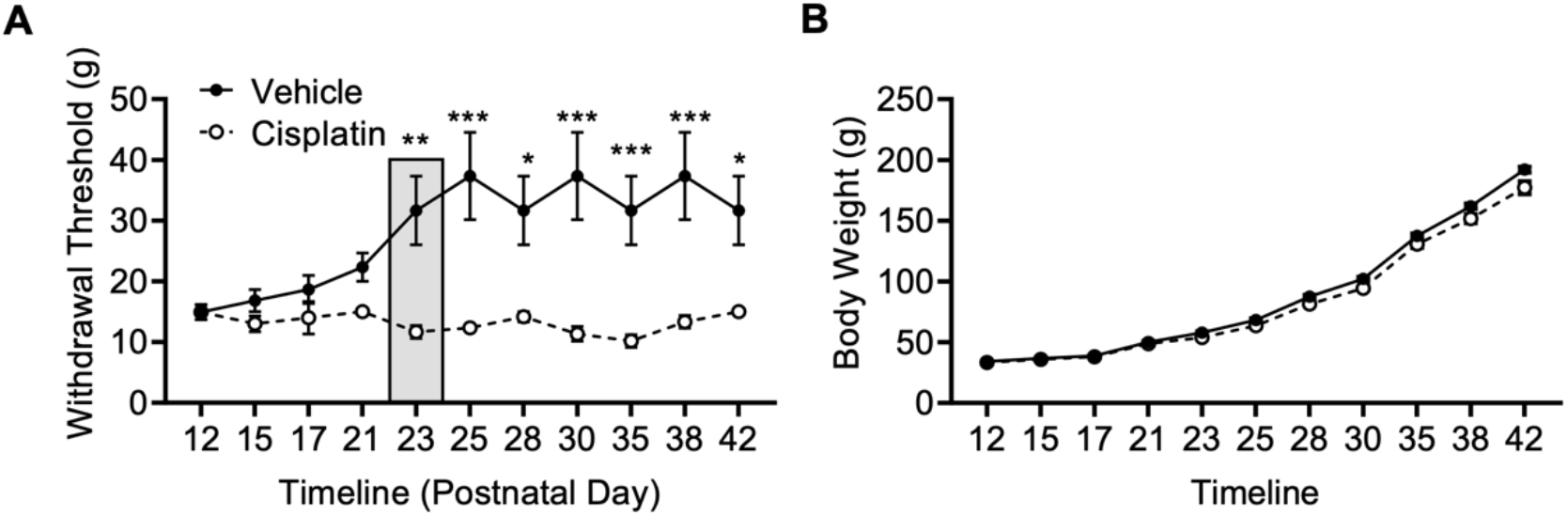
Early life exposure to cisplatin induces mechanical allodynia. [A] Neonatal exposure to cisplatin (IP 0.3mg/kg) led to a delayed (presenting at p23) but pronounced mechanical allodynia when compared to age matched sham control rodents. [B] There were no differences in body weight between vehicle and cisplatin treated rodents (Two way ANOVA with post Bonferroni, *p<0.05, **p<0.01, ***p<0.001).

**Figure 2.**
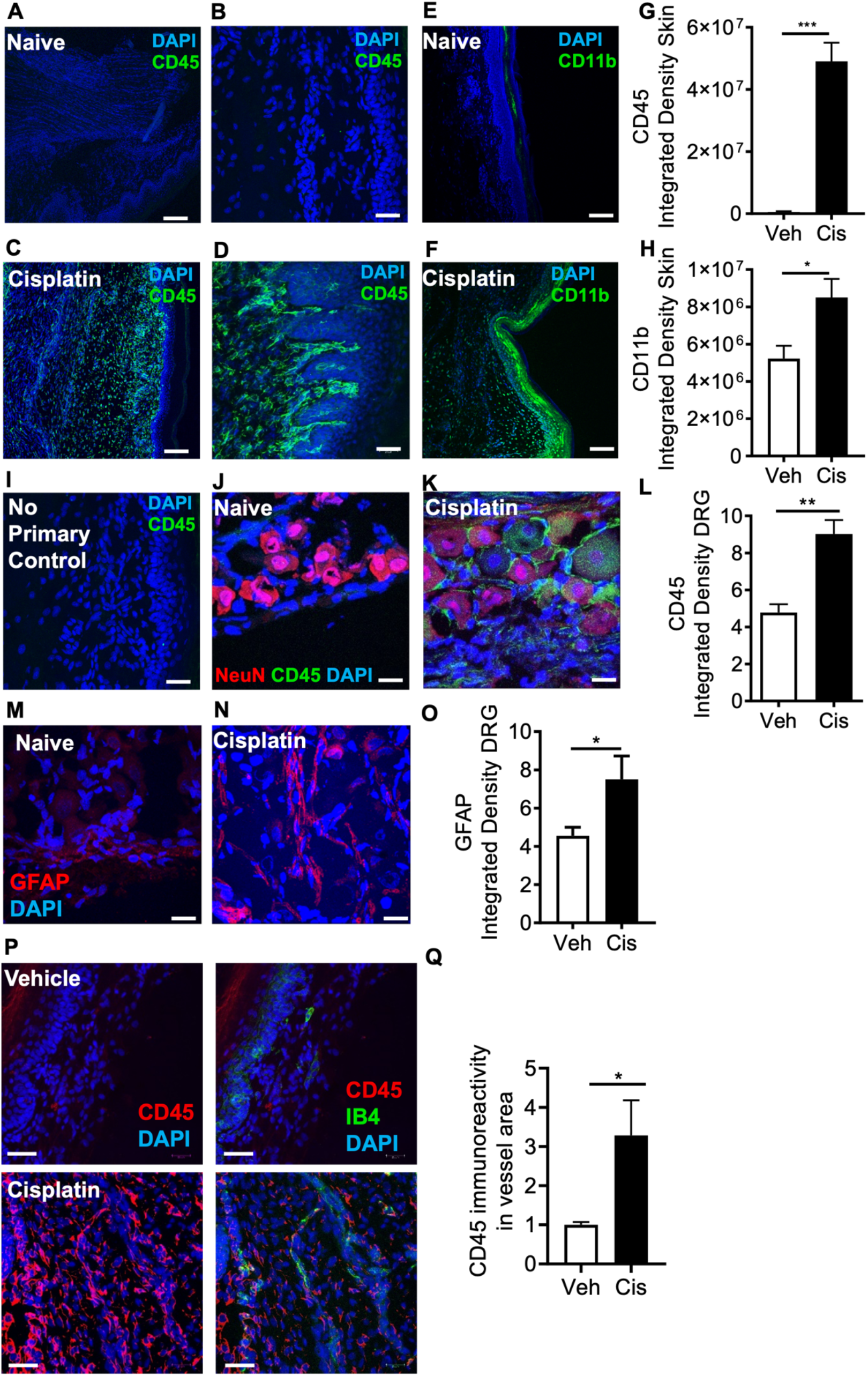
Cisplatin induced DRG macrophage infiltration. [A-D] Early life exposure of cisplatin led to an increase in proinflammatory environment in the peripheral sensory nervous system – dorsal root ganglia and plantar skin of the hindpaw. There was an increase in CD45 positive cells types in the plantar skin of the hindpaw in cisplatin treated rodents versus age matched sham control [G] (***p<0.001 Unpaired T Test, representative images [A&C] Scale bar = 50μm or [B&D] 25 μm). Similarly, [E&F] there was an increase in CD11b positive cells in the plantar skin of the hindpaw in cisplatin rodents versus age matched sham control rodents [H] (*p<0.05 Unpaired T Test, Scale bar = 50μm or 25 μm). Representative images of [I] no primary control. In [J] sham vehicle versus [K] cisplatin DRG demonstrated increased CD45 immunoreactivity. [L] In the dorsal root ganglia (lumbar 5) there was an increase in CD45 positive cells in cisplatin rodents when compared to age matched sham controls (*p<0.05 Unpaired T Test). Representative images of [M] sham vehicle versus [N] cisplatin DRG demonstrated GFAP immunoreactivity. [O] In the dorsal root ganglia there was an increase in GFAP positive cells in cisplatin rodents when compared to age matched sham controls. (*p<0.05 Unpaired T Test).

**Figure 3.**
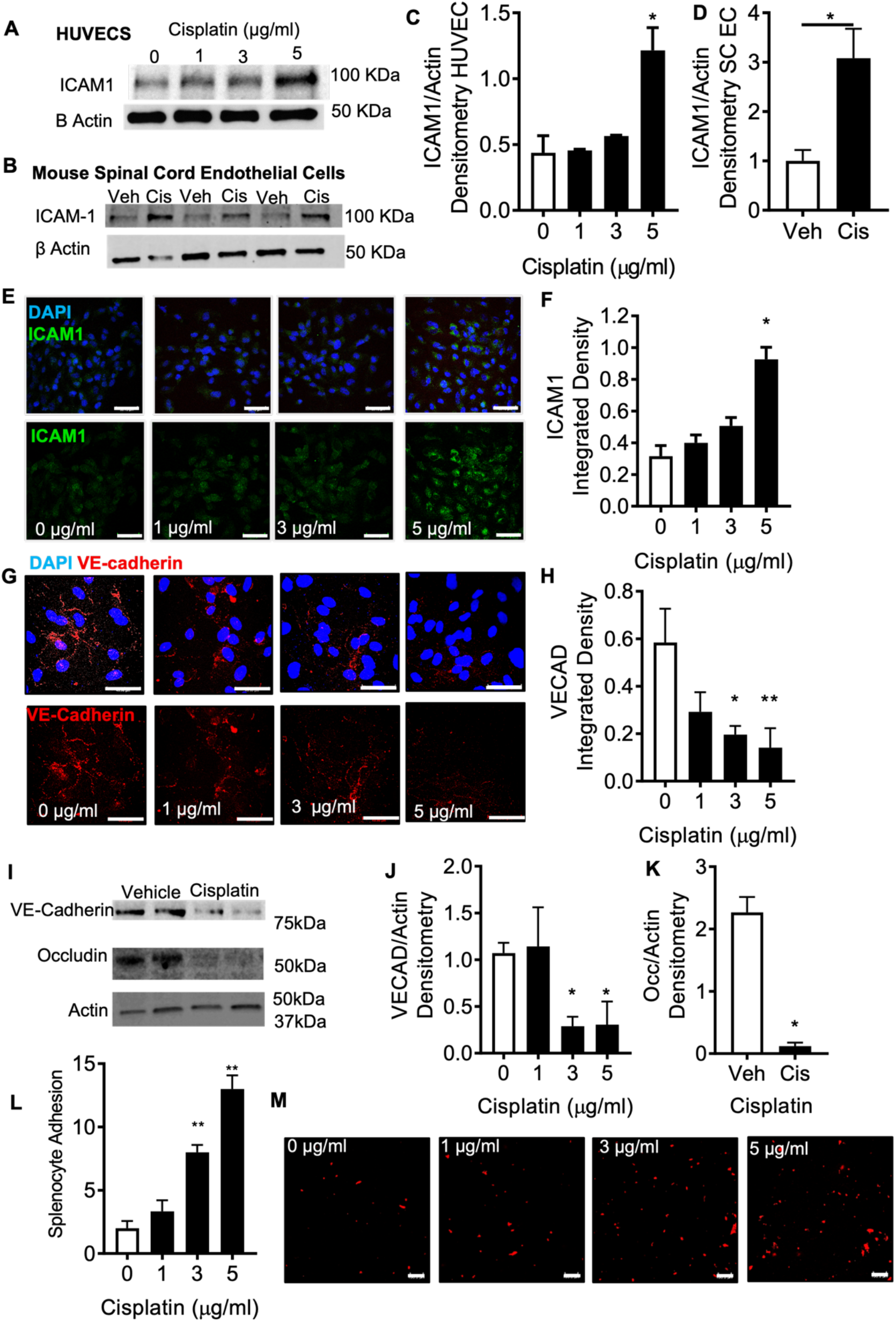
Cisplatin induced dorsal root ganglia inflammatory environment. Cisplatin-induced increases in ICAM-1 expression in [A,C] HUVECs and [B,D] mouse spinal cord endothelial cells as measured by densitometry analysis. Similarly using immunocytochemistry [E] ICAM1 expression was increased in HUVECS 24hrs post cisplatin treatment, [F] with increases occurring in a dose dependent manner (cisplatin concentrations 0, 1, 3 or 5 μg/ml). [G] HUVECs were treated with cisplatin led to reductions in endothelial tight junctional markers (VE Cadherin immunocytochemistry). [I] Protein extracted from HUVEC after 24h treatment with 0, 1, 3 or 5 μg/ml of cisplatin demonstrated reductions in [J] VE Cadherin and [K] Occludin expression. Immunocytochemistry of HUVECs treated for 24hrs with either vehicle or cisplatin demonstrated increased adherence of fluorescently labelled THP1 monocytes with increasing concentrations of cisplatin (One way ANOVA with post Bonferroni, *p<0.05, **p<0.01, Unpaired t Test *p<0.05, n= per group) Scale bar = 50μm). Cisplatin induced dorsal root ganglia vascular leakage and monocyte/macrophage infiltration. [A] In rodents treated with cisplatin early in life, there was an [B] increased accumulation of CD45 positive cells in the plantar skin that were associated with IB4 microvessels versus age matched control animals. (Unpaired t Test *p<0.05, scale bar = 25μm).

In the lumbar dorsal root ganglia extracted from cisplatin treated rodents there was increased accumulation of NGF positive cells (Fig. 4A&B) versus age matched controls. This accumulation of NGF in the DRG aligned with increased infiltrating CD45 positive monocyte/macrophage, with CD45 positive monocytes/macrophages expressing NGF (Fig. 4C&D). In isolated mouse splenocytes, cisplatin induced increased expression of NGF when compared vehicle treated splenocytes (Fig. 4E-F), as well as increasing the number of F480 (Fig. 4G) and CD11b (Fig. 4H). positive cells that express NGF post cisplatin treatment.

**Figure 4.**
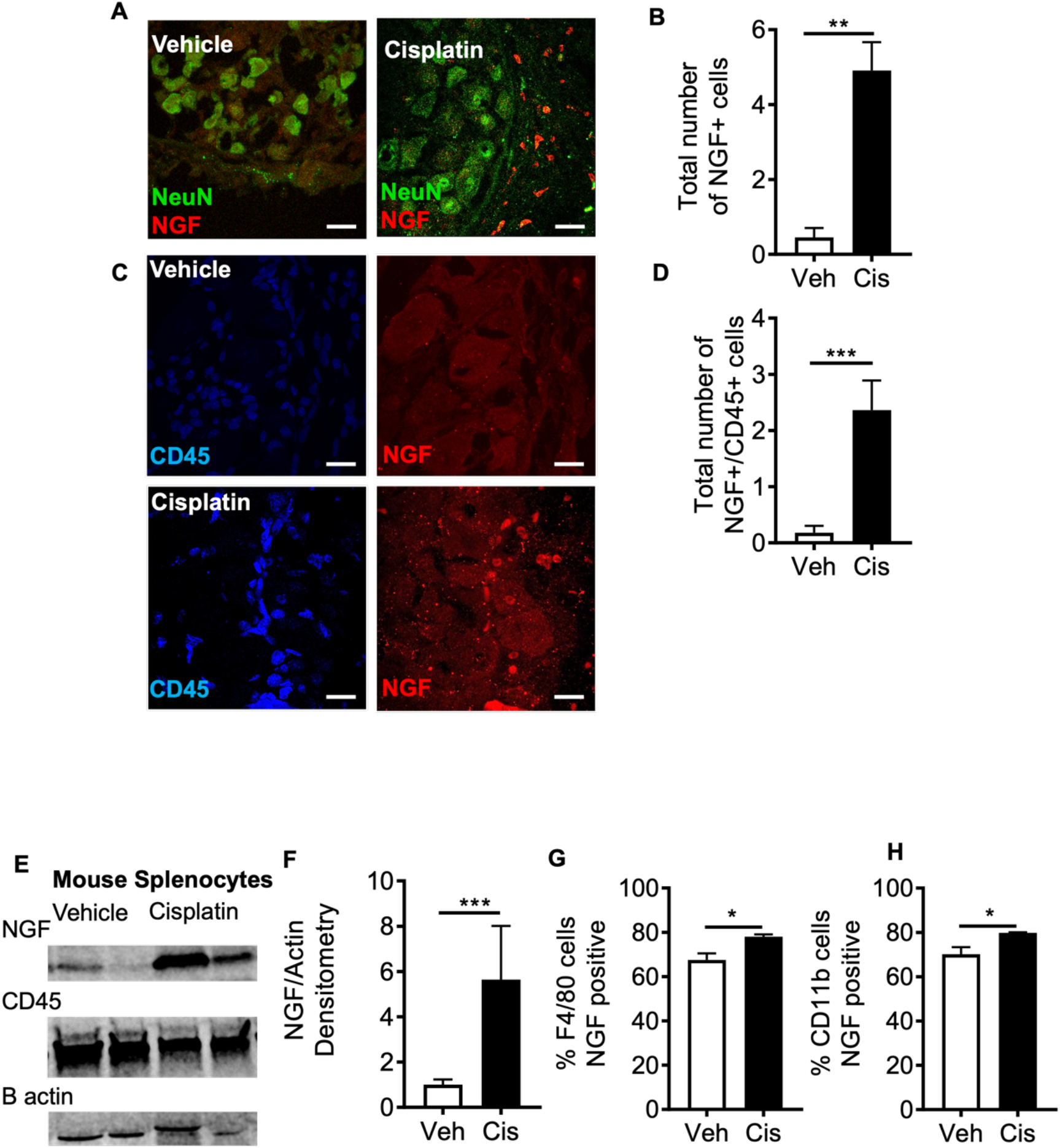
Cisplatin induced accumulation of NGF positive monocyte/macrophages in the dorsal root ganglia. [A] In rodents treated with cisplatin early in life, there was [B] increased number of NGF positive cells in the dorsal root ganglia (Scale bar = 25μm). Furthermore, [C&D] there was an increase in number of NGF positive CD45 cells in the DRG of cisplatin treated rodents versus sham vehicle control rodents (Scale bar = 25μm). [E] Isolated mouse splenocytes treated with either vehicle or cisplatin for 24hrs led to [E representative western blot &F] increased expression of NGF following cisplatin treatment and increased number of cells expressing NGF [G&H]. (Unpaired t Test *p<0.05, *** p<0.001).

NGF treatment led to increased capsaicin induced intracellular calcium influx (Fig.5A&B) in DRG nociceptors. Furthermore, cotreatment with neutralising NGF-antibody inhibited NGF induced DRG nociceptor sensitisation, whereas IgG control did not suppress NGF induced nociceptor sensitisation (Fig. 5A&B). Subcutaneous administration in the hindpaw plantar surface of NGF with neutralising NGF antibody inhibited NGF induced mechanical (Fig.5C) and heat (Fig.5D) hypersensitivity in the ipsilateral hindpaw. Subcutaneous administration of NGF into the plantar surface of the hindpaw led to increased number of CGRP positive intraepidermal nerve fibres (IENF) and branchpoints in the ipsilateral plantar skin (representative image CGRP positive IENF Fig.5E&G) versus vehicle treated group. NGF administered inconjunction with NGF antibody, inhibited NGF induced increased number of CGRP positive IENF and branchpoints (Fig.5E-G).

**Figure 5.**
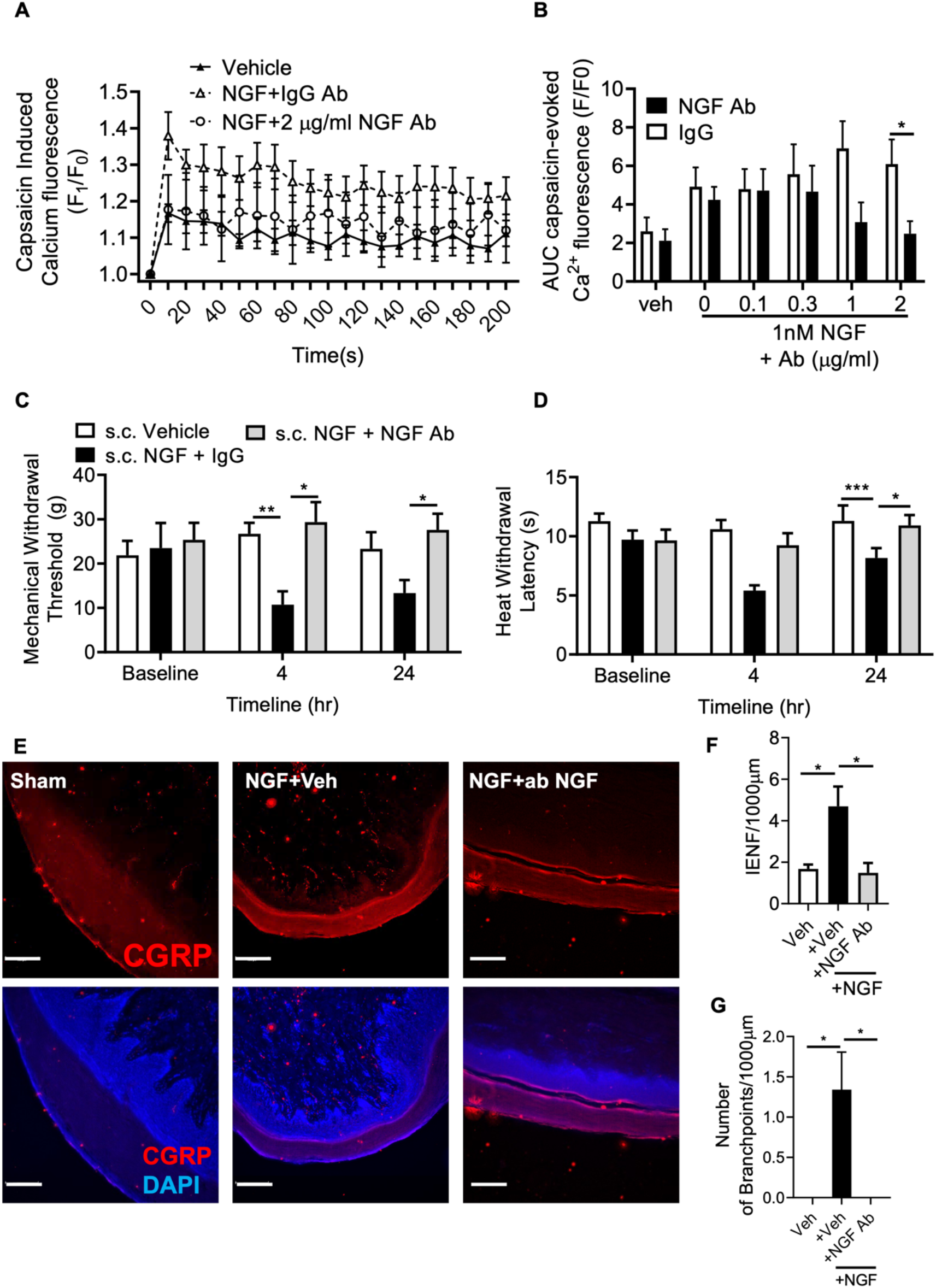
Neutralising antibody suppression in NGF induced nociceptor senstisation and aberrant growth. In isolated DRG primary sensory neuronal cell cultures, [A&B] NGF induced increase capsaicin mediated intracellular calcium influx, which was suppressed in a concentration dependent manner with increasing concentrations of NGF neutralising antibody (n=16 for PBS and NGF + 0μg/mL Ab treatment groups, n=14 for vehicle treatment, n=10 for NGF + 0.1μg/mL, 0.3μg/mL, 1μg/mL and 2μg/mL antibody treatment groups). Subcutaneous injection of NGF plus neutralising antibody preventing NGF induced [C] mechanical and [D] heat hyperalgesia, [E] alongside reductions in NGF induced CGRP nociceptor IENF [F] aberrant growth and [G] branch number versus NGF treatment. (One or Two way ANOVA with post Bonferroni, *p<0.05, **p<0.01, ***p<0.001). (Scale bar = 100μm).

DRG primary cell cultures were treated with cisplatin for 24hrs followed by a 7 day washout period. NGF induced TRPV1 sensitisation was exacerbated by cisplatin treatment, and treatment inconjunction with NGF neutralising antibody diminished NGF induced capsaicin responses compared to IgG control (Fig. 6A&B). Isolated mouse splenocytes were treated with either vehicle or cisplatin. Conditioned media was collected 24hrs later and applied to DRG primary cell cultures. Capsaicin induced intracellular calcium influx was increased in response to cisplatin treated conditioned media (Fig. 6C&D). Furthermore, with increasing concentrations of NGF neutralising antibody cisplatin treated conditioned media induced nociceptor sensitisation was inhibited (Fig.6C&D). In a rodent model of early life cisplatin induced chronic pain, mechanical allodynia was prevented following intraperitoneal injection of NGF neutralising antibody (Fig. 7A). In addition, intraperitoneal injection of NGF neutralising antibody prevented cisplatin induced mechanical allodynia in male (Fig. 7C) and female (Fig. 7D) rodents. Furthermore, cisplatin induced increases in CGRP positive nociceptor IENF in the plantar skin was prevented in the NGF neutralising antibody treated group (Fig. 8B-C). In addition, cisplatin induced accumulation of CD45 positive cells in the plantar hindpaw skin was not altered by administration of NGF neutralising antibody (Fig. 8D&E).

**Figure 6.**
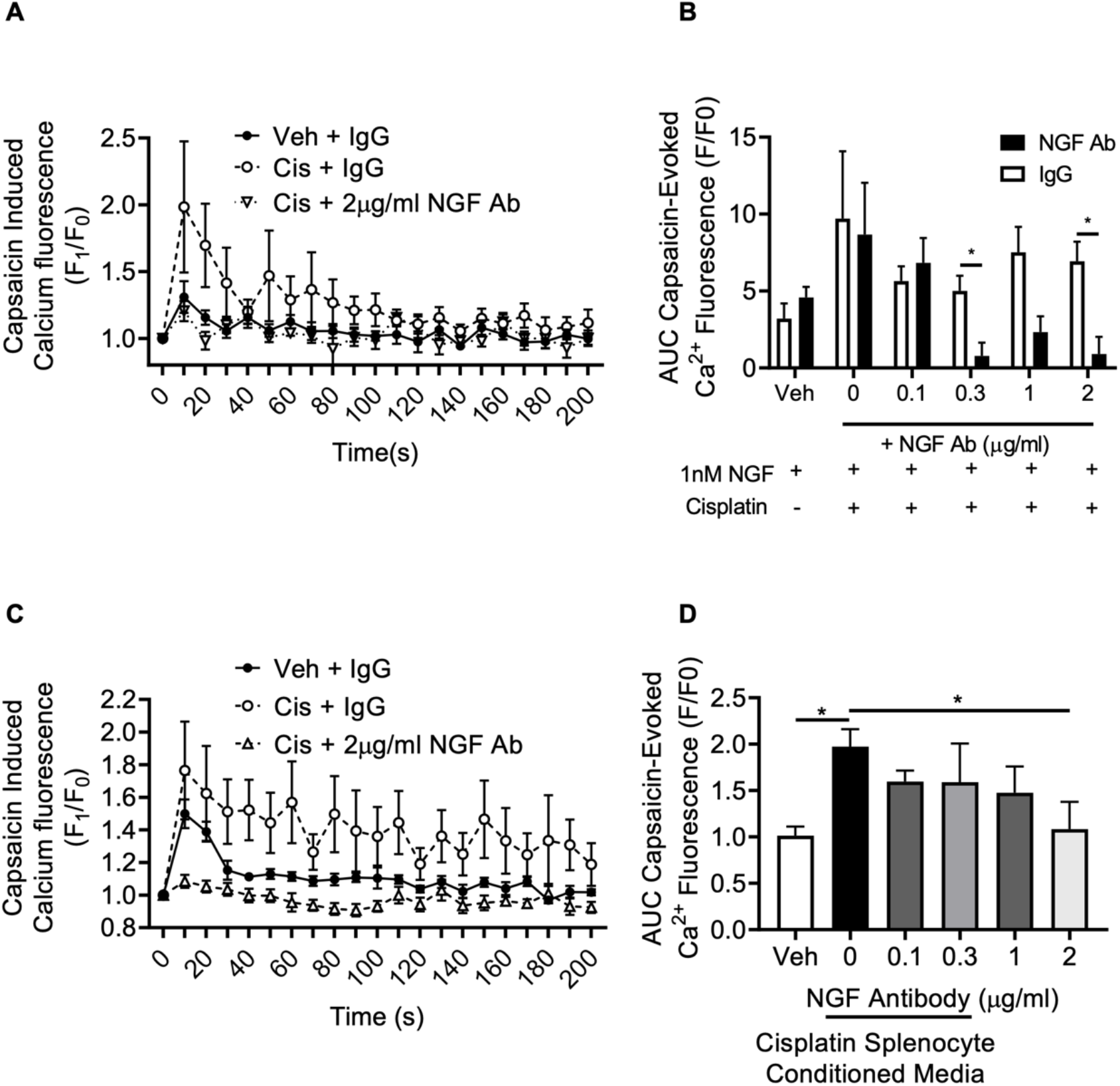
Cisplatin induced proinflammatory mediated nociceptor sensitisation is NGF dependent. [A&B] Treatment of with 5μg/mL cisplatin 1 week prior to NGF treatment increased TRPV1 activity when compared to vehicle. NGF neutralising antibody diminished capsaicin induced TRPV1 mediated nociceptor activity. [C&D] Conditioned media from cisplatin treated mouse splenocytes led to increased capsaicin evoked DRG nociceptor activity compared vehicle treated conditioned media from mouse splenocytes, which was diminished with increasing concentrations NGF neutralising antibody. (One way ANOVA with post Bonferroni, *p<0.05).

**Figure 7.**
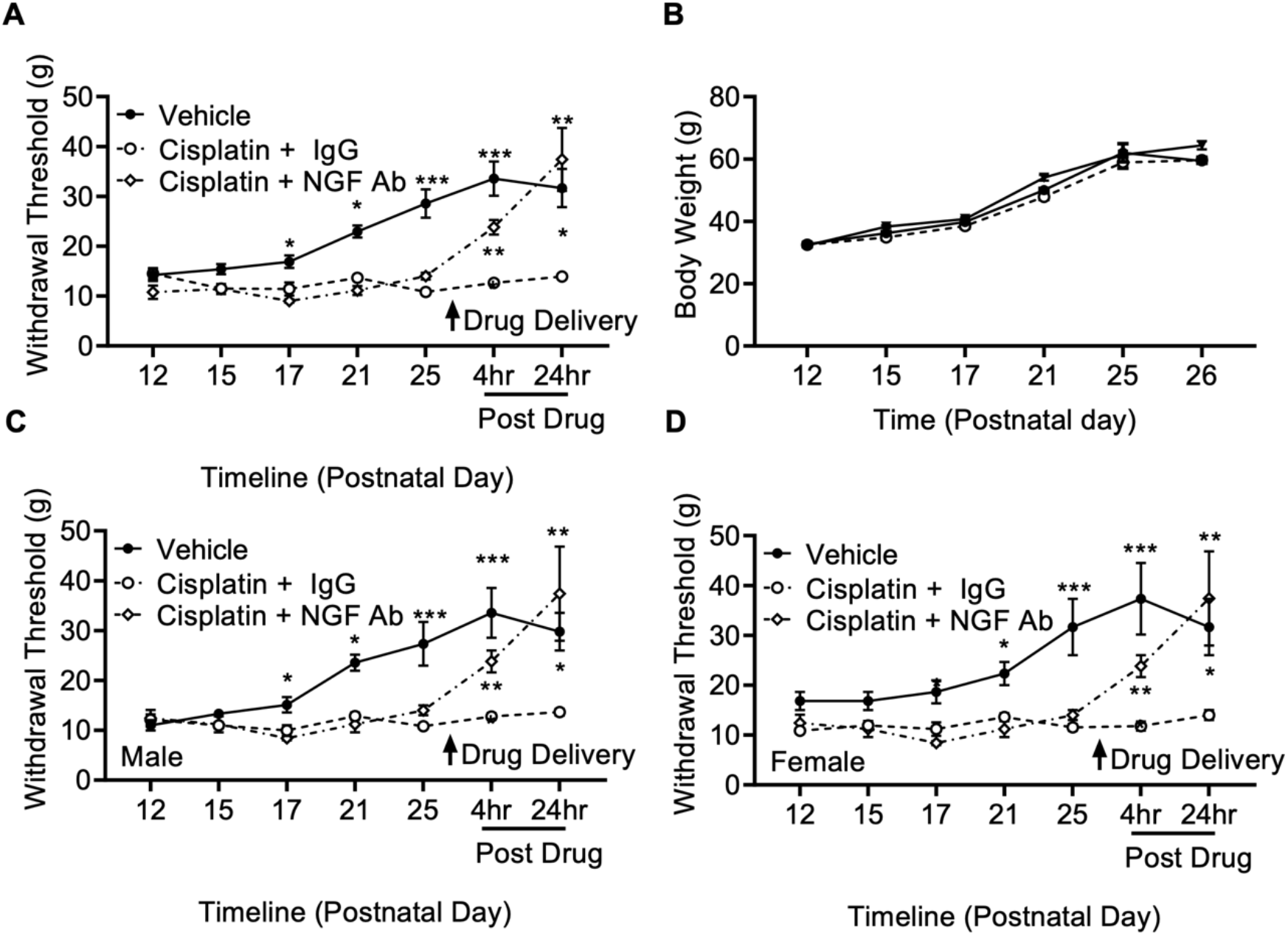
NGF mediated cisplatin induced survivorship pain. In a rodent model of early life exposure of cisplatin induced chronic pain [A] intraperitoneal injection of NGF neutralising antibody prevented mechanical allodynia versus cisplatin + vehicle group. [B] Body weight did not differ between each experimental group. In addition, NGF neutralising antibody prevented mechanical allodynia versus cisplatin + vehicle group in [C] males and [D] female rodents. (Two way ANOVA with post Bonferroni, *p<0.05, **p<0.01, ***p<0.001).

**Figure 8.**
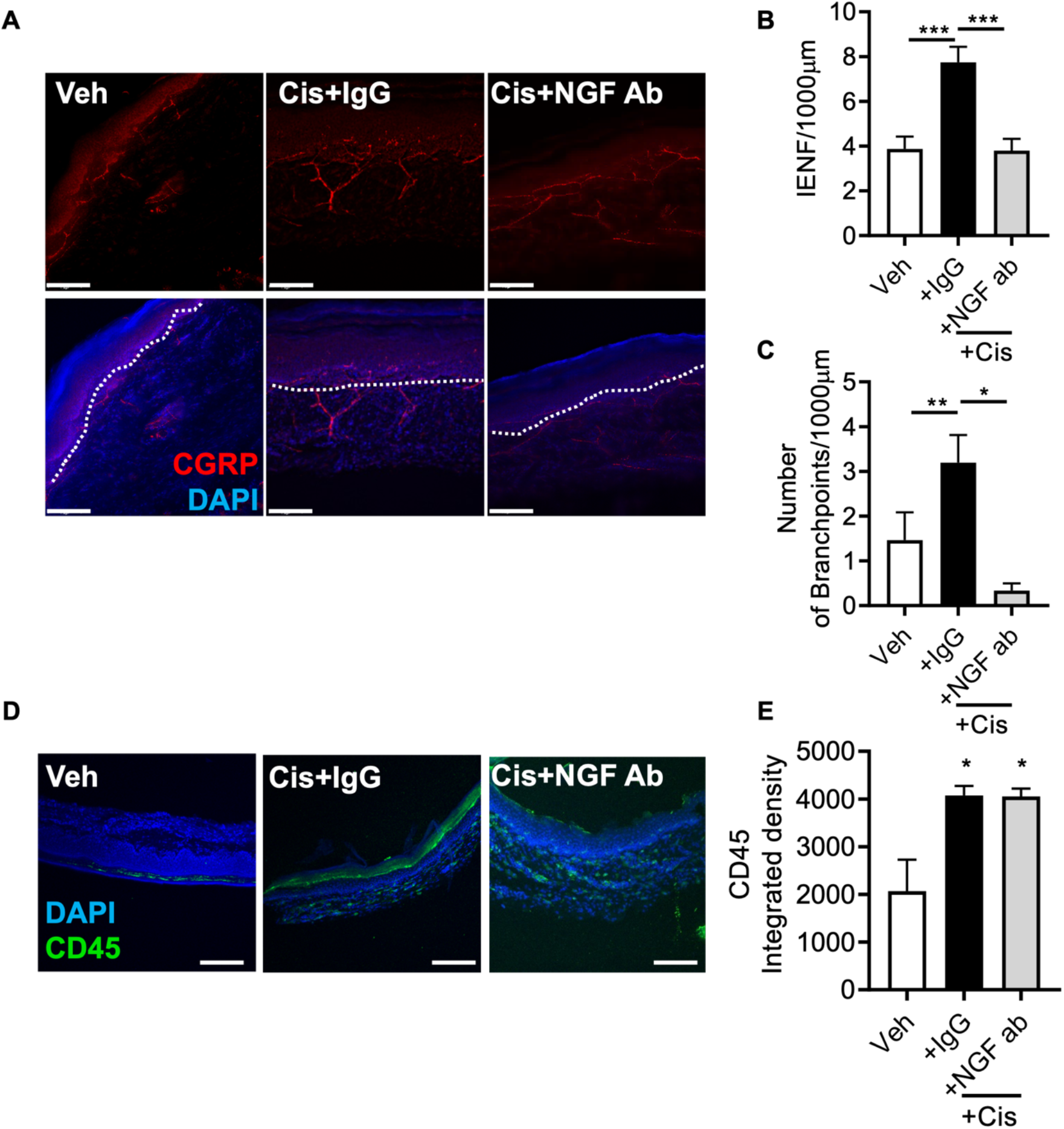
Cisplatin induced aberrant IENF growth is NGF dependent. [A representative pictures] Early life exposure to cisplatin led to [B] increased CGRP positive nociceptor IENF in the plantar skin versus sham vehicle group CGRP nociceptor and [C] increased IENF branch number. In addition, NGF neutralising antibody prevented cisplatin induced aberrant growth and branch number in CGRP positive IENF. [D] Cisplatin induced a proinflammatory environment (increased number of CD45 positive cells) in the plantar skin versus vehicle sham treated group. [E] Proinflammatory state (CD45 positive cell infiltration] within the plantar skin remained unchanged between cisplatin and cisplatin + NGF antibody. (One way ANOVA with post Bonferroni, *p<0.05, **p<0.01, ***p<0.001). (Scale bar = 100μm).

## Discussion

In this study we demonstrate that early life treatment with cisplatin instigates a prominent neuroinflammatory component in the delayed but lasting manifestation of neuropathic pain states, which was comparable between male and female mice. Cisplatin induced pain and nociceptor sensitisation was driven by infiltrating M1 like macrophages into the peripheral somatosensory nervous system (PNS) in an NGF dependent. This investigation outlines an inflammatory mechanism that induces the onset of pain in adult survivors of childhood cancer.

### Neuroinflammatory induced childhood cancer survivorship pain

A large proportion of adult individuals whom have been treated during infancy for paediatric cancer, present a significant neuropathic pain phenotype, lasting into adulthood. Current analgesic therapies used to treat platinum-based neuropathic pain provides minimal effective pain relief [2]. Current understanding of neuropathic pain development following early life exposure to chemotherapeutic agents demonstrates that pain hypersensitivity is delayed, presenting during adolescence. Furthermore, there is increasing presentation occurring with earlier exposure to chemotherapeutic agents as well as pain symptoms increasingly apparent with advancing age. Current clinical studies present the development of a lasting pain in adult paediatric cancer patients, however, the ongoing underlying mechanisms that cause this sensory neuropathology has not been elucidated. Recently, several rodent models have been developed to enable exploration of the etiology of this complication and to expand our understanding. It is widely regarded that chemotherapy induced neuropathic pain is dependent upon sensory neurotoxicity due to the nature of platinum based agents accumulating in the PNS [4], inducing nociceptor sensitisation. Current understanding of developmental sensory neurobiology is that the delay in the presentation of neuropathic pain phenotypes until adolescence following early life exposure to a ‘neural stressor’ aligns with the development of proinflammatory processes. Exposure of the developing immature PNS to a traumatic/toxic insult initiates maladaptations within the developmental trajectory of the nociceptor and surrounding environment to drive a delayed but lasting chronic pain phenotype in humans and rodents. Part of this is the unsilencing of the immune system during adolescence, which is normally during neural development anti-inflammatory in nature preventing nociceptor activation. During adolescence, early life exposure to cisplatin initiates in infancy alterations in the vascular-immune interactions. Promoting macrophage infiltration into the PNS through increased capillary endothelium permeability that aligns with enhanced macrophage adhesion, trafficking and penetration. Furthermore, infiltrating macrophages initiate a DRG microenvironment that is abundant with nociceptor sensitising agents/inflammatory mediators. It has previously been shown in other complications that platinum based chemotherapy drives inflammation in a multitude of systems, for instance in cisplatin induced nephropathy, whereby macrophage infiltration is associated with pathology progression. Exposing nociceptors to the secretome of cisplatin treated macrophages initiates nociceptor sensitisation. This is particularly pertinent as cisplatin induced survivorship pain is associated with an increased infiltration of CD45 cells and GFAP immunoreactivity restricted to the DRG; depicting neuroinflammatory activation.

### NGF dependent induction childhood cancer survivorship pain

Chemotherapy induces neuropathic pain in adults, however in infants the onset of exaggerated pain phenotypes does not manifest until later in life. Previous investigation has identified alterations in the maturation of the developing nociceptor following early exposure to platinum based chemotherapy, cisplatin. Cisplatin induced survivorship pain has been associated with aberrant nociceptor growth in the peripheral tissues, in this instance skin. Exaggerated abnormal intraepidermal nerve fibre growth with particular reference to calcitonin gene related peptide (CGRP) labelled nociceptors is widely established to align with onset of chronic pain states, typically in times of tissue inflammation that occurs during cancer and arthritis. Inflammation and the local inflamed tissue environment promotes hyperalgesia and tissue sensitisation. Key to the efficient function of the PNS is the supporting glial and inflammatory cell types, which secrete cytokines such as nerve growth factor (NGF). Activation of these cell types (neuroinflammation) is crucial to the development of chronic pain in adults as well as in children. Inhibition of inflammatory signalling such as NGF and TNF alpha mediated pathways are currently targeted for analgesic development in arthritis. There are numerous factors that influence developmental trajectory of nociceptor skin innervation. A primary modulator is nerve growth factor (NGF). NGF coordinates and establishes IENF during development, with waves of elevated NGF expression occuring during development of sensory afferent terminals and with IENF hardwiring becoming established at 3 weeks of life. However, surgical intervention and induction of local tissue inflammation within first week of life induces elevations in NGF expression concomitantly is associated with regenerative/neurogenesis of sensory afferents terminals through elevated levels of growth associated protein 43 (GAP43) and increases in nociceptor skin innervation. In addition, NGF is associated with nociceptor mediated hyperalgesia and long term nociceptor sensitisation demonstrated by hyperalgesic priming.

The immune system plays a significant role in the mechanisms behind CIPN manifestation; cisplatin has been found to activate and recruit inflammatory cells to the area that release pro-inflammatory cytokines, such as chemokines (CKs) and interleukins (ILs). Studies have found that elevated levels of IL-1b, IL-6, tumour necrosis factor-a (TNF-a) as part of the inflammatory response associated with CIPN. Here we present NGF as a the first mediator of cisplatin induced survivorship pain due to the induction of M1 polarisation. The upregulation of these molecules is thought to stimulate the hyperexcitability and sensitisation of peripheral nociceptive neurons that cause the perceived pain sensation in CIPN patients. The dorsal root ganglia are particularly vulnerable to inflammatory damage due to a highly permeable protective barrier [10]. Infiltration of differing inflammatory cell types promotes a proinflammatory environment within the dorsal root ganglia of cisplatin treated rodents. There have been extensive studies regarding peripheral sensitisation by varying inflammatory mediators in response to injury, including the role of NGF/TrkA signalling in the pathophysiology of peripheral sensory neuropathy. As NGF/TrkA signalling has been established as a key mediator in the development of neuropathic pain, inhibition of this pathway appears as an attractive target as a novel analgesic therapy. Hathway and colleagues investigated the effects of cisplatin treatment on the maturation of nociceptive systems in an in vivo rodent model and found an increase in TrkA+ neurons in the cisplatin-treated group [9]. Importantly, patients treated with cisplatin have been reported as having elevated serum NGF concentrations [21] highlighting that NGF-TrkA signalling axis as a prominent mediator of cisplatin induced neuropathic pain.

Our findings establish that NGF dependent modulation of cisplatin induced DRG nociceptor sensitisation and consequent development of chronic pain states. The results from this study indicate that cisplatin treatment exacerbates NGF mediated nociceptor sensitisation, and therefore acts as a key mediator in increased pain perception in chemotherapy-induced pain in adult survivors of childhood cancer. Furthermore, the present findings demonstrate the clinical relevance of developing an antibody-based novel analgesic to inhibit NGF mediated nociceptor signalling, in order to treat the painful symptoms of chemotherapy induced peripheral neuropathy in childhood survivors of cancer.

## Acknowledgements

RPH, LH, TV, MD, LT, CG, DS, JE and MPC performed the experimental work and contributed to the conception or design of the work in addition to acquisition, analysis or interpretation of data for the work. All authors drafted the article or revised it critically for important intellectual content. All authors approved the final version of the manuscript.

This work was supported by the European Foundation for the Study of Diabetes Microvascular Programme supported by Novartis to RPH (Nov 2015_2 to RPH), the EFSD/Boehringer Ingelheim European Research Programme in Microvascular Complications of Diabetes (BI18_5 to RPH), the Rosetree Trust (A1360 to RPH) and Nottingham Trent University.

The authors would like to thank Graham Hickman of the Imaging Suite at Nottingham Trent University for their support and assistance in this work.

## References

[1] Alberts NM, Gagnon MM, Stinson JN. Chronic pain in survivors of childhood cancer: a developmental model of pain across the cancer trajectory. Pain 2018. doi:10.1097/j.pain.0000000000001261.

[2] Argyriou AA, Koutras A, Polychronopoulos P, Papapetropoulos S, Iconomou G, Katsoulas G, Makatsoris T, Kalofonos HP, Chroni E. The impact of paclitaxel or cisplatin-based chemotherapy on sympathetic skin response: A prospective study. Eur J Neurol 2005;12:858–861.

[3] Drake RA, Steel KA, Apps R, Lumb BM, Pickering AE. Loss of cortical control over the descending pain modulatory system determines the development of the neuropathic pain state in rats. Elife 2021;10.

[4] Ferdousi M, Azmi S, Petropoulos IN, Fadavi H, Ponirakis G, Marshall A, Tavakoli M, Malik I, Mansoor W, Malik RA. Corneal confocal microscopy detects small fibre neuropathy in patients with upper gastrointestinal cancer and nerve regeneration in chemotherapy induced peripheral neuropathy. PLoS One 2015;10.

[5] Fitzgerald M. The development of nociceptive circuits. Nat Rev Neurosci 2005;6:507–520. doi:10.1038/nrn1701.

[6] Fitzgerald M, McKelvey R. Nerve injury and neuropathic pain - A question of age. Exp Neurol 2016;275 Pt 2:296–302. doi:10.1016/j.expneurol.2015.07.013.

[7] Gilchrist LS, Tanner L. The pediatric-modified total neuropathy score: a reliable and valid measure of chemotherapy-induced peripheral neuropathy in children with non-CNS cancers. Support Care Cancer 2013;21:847–856. doi:10.1007/s00520-012-1591-8.

[8] Hargreaves K, Dubner R, Brown F, Flores C, Joris J. A new and sensitive method for measuring thermal nociception in cutaneous hyperalgesia. Pain 1988;32:77–88.

[9] Hathway GJ, Murphy E, Lloyd J, Greenspon C, Hulse RP. Cancer Chemotherapy in Early Life Significantly Alters the Maturation of Pain Processing. Neuroscience 2017.

[10] Jimenez-Andrade JM, Herrera MB, Ghilardi JR, Vardanyan M, Melemedjian OK, Mantyh PW. Vascularization of the dorsal root ganglia and peripheral nerve of the mouse: Implications for chemical-induced peripheral sensory neuropathies. Mol Pain 2008;4:1–8.

[11] Khan RB, Hudson MM, Ledet DS, Morris EB, Pui CH, Howard SC, Krull KR, Hinds PS, Crom D, Browne E, Zhu L, Rai S, Srivastava D, Ness KK. Neurologic morbidity and quality of life in survivors of childhood acute lymphoblastic leukemia: a prospective cross-sectional study. J Cancer Surviv 2014;8:688–696.

[12] Lu Q, Krull KR, Leisenring W, Owen JE, Kawashima T, Tsao JC, Zebrack B, Mertens A, Armstrong GT, Stovall M, Robison LL, Zeltzer LK. Pain in long-term adult survivors of childhood cancers and their siblings: a report from the Childhood Cancer Survivor Study. Pain 2011;152:2616–2624. doi:10.1016/j.pain.2011.08.006.

[13] McKelvey R, Berta T, Old E, Ji RR, Fitzgerald M. Neuropathic pain is constitutively suppressed in early life by anti-inflammatory neuroimmune regulation. J Neurosci 2015;35:457–466.

[14] Ness KK, Jones KE, Smith WA, Spunt SL, Wilson CL, Armstrong GT, Srivastava DK, Robison LL, Hudson MM, Gurney JG. Chemotherapy-related neuropathic symptoms and functional impairment in adult survivors of extracranial solid tumors of childhood: results from the St. Jude Lifetime Cohort Study. Arch Phys Med Rehabil 2013;94:1451– 1457.

[15] Phillips SM, Padgett LS, Leisenring WM, Stratton KK, Bishop K, Krull KR, Alfano CM, Gibson TM, de Moor JS, Hartigan DB, Armstrong GT, Robison LL, Rowland JH, Oeffinger KC, Mariotto AB. Survivors of Childhood Cancer in the United States: Prevalence and Burden of Morbidity. Cancer Epidemiology Biomarkers & Prevention 2015;24:653–663. doi:10.1158/1055-9965.epi-14-1418.

[16] Robison LL, Hudson MM. Survivors of childhood and adolescent cancer: Life-long risks and responsibilities. Nat Rev Cancer 2014;14:61–70.

[17] Schindelin J, Arganda-Carreras I, Frise E, Kaynig V, Longair M, Pietzsch T, Preibisch S, Rueden C, Saalfeld S, Schmid B, Tinevez JY, White DJ, Hartenstein V, Eliceiri K, Tomancak P, Cardona A. Fiji: An open-source platform for biological-image analysis. Nat Methods 2012;9:676–682. doi:10.1038/nmeth.2019.

[18] Schneider CA, Rasband WS, Eliceiri KW. NIH Image to ImageJ: 25 years of image analysis. Nat Methods 2012;9:671–675.

[19] Schwaller F, Fitzgerald M. The consequences of pain in early life: injury-induced plasticity in developing pain pathways. Eur J Neurosci 2014;39:344–352. doi:10.1111/ejn.12414.

[20] Ved N, Lobo MEDV, Bestall SM, Vidueira CL, Beazley-Long N, Ballmer-Hofer K, Hirashima M, Bates DO, Donaldson LF, Hulse RP. Diabetes-induced microvascular complications at the level of the spinal cord; a contributing factor in diabetic neuropathic pain. J Physiol 2018;596:3675–3693. doi:10.1113/JP275067.

[21] Velasco R, Navarro X, Gil-Gil M, Herrando-Grabulosa M, Calls A, Bruna J. Neuropathic Pain and Nerve Growth Factor in Chemotherapy-Induced Peripheral Neuropathy: Prospective Clinical-Pathological Study. J Pain Symptom Manage 2017;54:815–825.

[22] Da Vitoria Lobo ME, Weir N, Hardowar L, Al Ojaimi Y, Madden R, Gibson A, Bestall SM, Hirashima M, Schaffer CB, Donaldson LF, Bates DO, Hulse RP. Hypoxia induced carbonic anhydrase mediated dorsal horn neuron activation and induction of neuropathic pain. Pain 2022;163:2264–2279.

